# Social interaction influences innate color preference of zebrafish shoals

**DOI:** 10.1101/2020.03.23.003186

**Authors:** Ju Wang, Lifen Yin, Bin Hu, Lei Zheng

## Abstract

The color discrimination can confer survival advantages by helping animals to find nutritious food and shelter and to avoid predator. Zebrafish as a social species, data on innate color preference in shoals remain controversial and there are limited data for this organism. Here we showed that, when given a choice among two color combinations (R-Y, R-G, Y-G, B-G, B-R, B-Y), shoals of zebrafish exhibited a complex pattern of color preference and the order of RYGB preference was R>Y>G, B>G. By contrast, the individual zebrafish showed marked changes, completely losing their preference for all the tested color combinations. To investigate the role of shoaling behavior in color preference, we selected a D1-receptor antagonist (SCH23390), which could disrupt social preference and decrease social interaction in zebrafish. Interestingly, the shoals that were treated by SCH23390 showed no color preference for all color combinations. Our findings indicate that social interaction is involved in color-driven behavior in zebrafish, and reveal the possible mechanisms that the dopaminergic system may contribute to innate color preference in shoals of zebrafish.

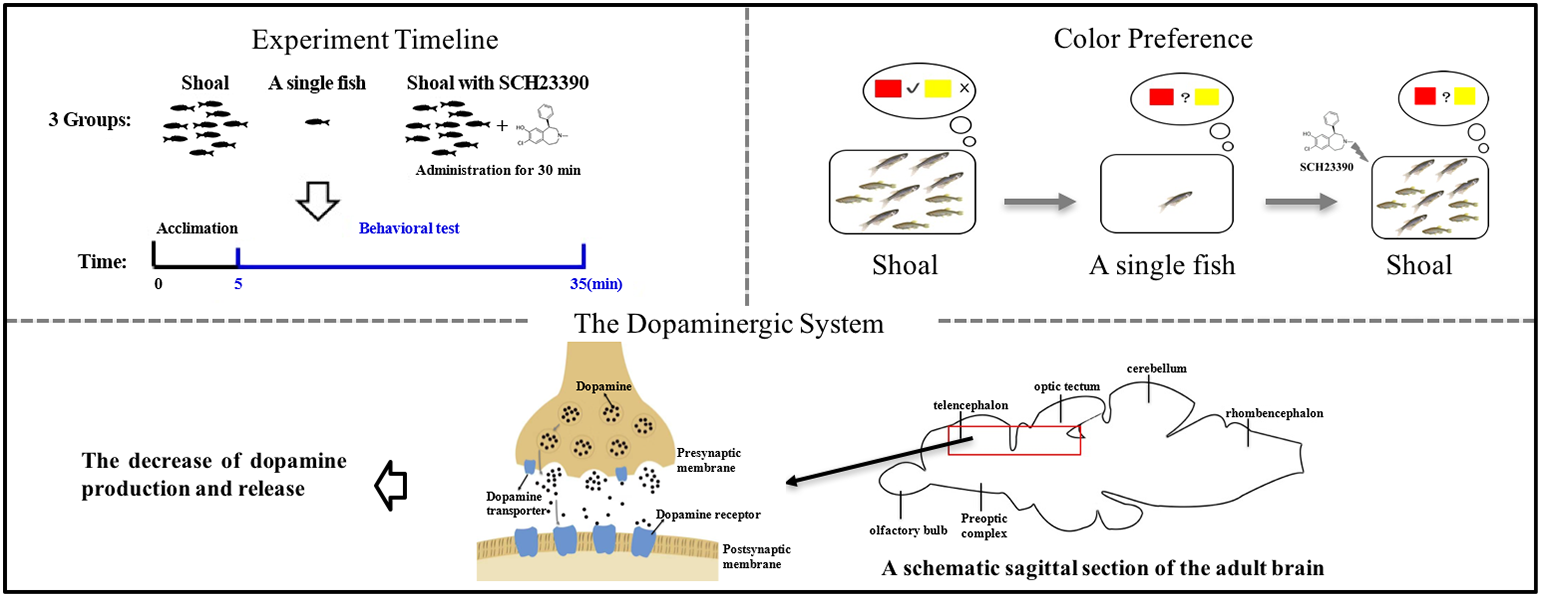
Graphical Abstract.

## 1. Introduction

Color preference has been studied in a wide range of species-insects [1–3], birds [4], fish [5–7], and humans [8]- affecting foraging, decision making, and reproduction. Humans are commonly the most like blue and dislike yellowy-green when individuals vary in their color preference [8]. Animals are often better able to perceive colors in their environment [9–11], and their color preference are either most represented in the environment [12] or contrast with the background [13]. For example, the parasitic wasp (*Venturia canescens*) prefers yellow, which is the most common color among natural flowers in their living regions [1]. Bumblebees (*Bombus impatiens*) prefer blue flowers, which stand out in a complex background [2]. The fruit fly *Drosophila* prefers green light to red light in the early morning and late afternoon, when the flies showed the higher activity, because such timed preference and the burst of activity are devoted to searching for food in or under green trees and bushes [3,14]. While tropical Asian birds prefer red and black, which are the most commonly encountered forest fruit colors [4].

Plenty of researches have proven that zebrafish (*Danio rerio*) have preference towards different colors [5–7, 15–17]. Surprising, the color preference of zebrafish has been extensively studied but still remains controversial. For instance, some researches show a strong preference for blue [5,15], whereas others report a clear aversion for this color [6,7,16,17]. It is still not clear whether the shoaling behavior affect the color preference. As is well known, zebrafish are social animals and tend to travel in shoals [18]. Meanwhile, there is a vital necessity for fish to sense danger and stay away from predators, thus it is beneficial for a fish to stay tight within a shoal and to assess the social interaction efficiently. We make a hypothesis that the characteristic of background color may influence the efficiency of social contact within a shoal and consequently shoal may prefer certain background colors in order to keep the social contact, which can be called as innate color preference of shoals. Recently, Park *et al*. used zebrafish larvae (5 days post fertilization (dpf)) to test the innate color preference in shoals [19]. However, shoaling behavior usually starts to develop after 7 dpf, becoming progressively stronger for the mature [20–22]. As far as we know, no research has been published that investigate the innate color preference of mature shoals, although shoaling and social behavior in general has received considerable critical attentions.

The cohesion of shoals has been found to be associated with the whole brain dopamine level [23]. Dopamine is one of the major neurotransmitters in the central nervous system of the vertebrate brain which plays important roles in a variety of cerebral functions, such as mood, attention, reward and memory [24–26]. Abundant evidence shows that dopamine is associated with the neurobehavioral functions in zebrafish [27,28]. Saif *et al*. [27] found that strong social stimuli will increase the dopamine and its metabolite 3,4-dihydroxyphenylacetic acid (DOPAC) levels in the brain of the adult zebrafish. The short-term isolated zebrafish could reduce the level of DOPAC [28]. The social interaction of shoals in zebrafish can be affected by the dopaminergic system through influence of the dopamine level [29]. D1 dopamine receptor antagonist (SCH23390) is most abundantly expressed dopamine receptor subtypes in the brain of zebrafish [30]. And SCH23390 disrupts social preference of zebrafish by decreasing the level of dopamine in dopaminergic system [31,32].

In the present study, we used two-color combinations (R-Y, R-G, Y-G, B-G, B-R, B-Y) to test the innate color preference of shoal (10 adult zebrafish) and individual fish, respectively. Moreover, we evaluated the influence of social interaction on the color preference of shoal, by implementing D1-receptor antagonist to disrupt social preference of zebrafish.

## 2. Materials and Methods

### 2.1. Animals

Adult zebrafish of the wild type (AB strain) were obtained from breeding center at University of Science and Technology of China. Zebrafish were maintained in an environmental controlled room with a 14 h light/10 h dark cycle (room fluorescent light, 08:00 am-22:00 pm) and a temperature at 28°C. The pH and conductivity in circulating water of the aquarium were 7.0-7.4 and 1500-1600 μs/cm, respectively. Animals were fed twice per day, at 09:00 am and 14:00 pm, with frozen brine shrimps.

### 2.2. SCH23390 exposure

1.0 mg/L D1-receptor antagonist SCH23390 (R-(+)-8-chloro-2,3,4,5-tetrahydro-3-methyl-5-phenyl-1H-3-benzazepine-1-ol; Cat # D054; Sigma-Aldrich) were applied to treat individuals. 1.0 mg/L SCH23390 which significantly reduced the amount of dopamine in the brain of zebrafish [29], so we selected this concentration to conduct our experiments. Before color preference test, fish were placed in drug exposure beaker (500 mL in volume) and remained in D1-receptor antagonist solution for 30 min. Because the 30-min exposure period is sufficiently long for the drug to reach the zebrafish brain through the vasculature of their gills and skin [29]. All the animals in the beaker were offered the same conditions (including illumination, temperature and dissolved oxygen) which were identical to the standard aquarium.

### 2.3. Experimental apparatus

The color-enriched conditional place preference (CPP) apparatus, is a commercial fish tank (35 cm length × 20 cm width × 23 cm height), colored with four color combinations (red (R), green (G), yellow (Y) and blue (B)). To create the preference for two colors, the CPP tank was divided into two compartments which were covered with the corresponding colors on all side except the top. A video camera (NVH-589MW; Wang Shi Wu You Corporation; China) was placed above the CPP tank for vertical video tracking. The color preference apparatus was placed over the LED light panel to ensure the light source could cover the whole tank. The detailed apparatus has been described in our previous work [33]. The experimental tanks were poured into 10 L fresh fish water, there was no physical barrier between the two compartments and the experimental subjects could swim freely in the entire tank. The outside of color-enriched CPP tank was opaque to prevent external visual interference from all direction. To minimize the effect of noise, the experimental room was closed and kept quiet, and the experimenter was not visible to the fish during the recording.

### 2.4. Color preference test

To test the color preference of shoals in zebrafish, the color-enriched CPP tank is for to measurement of color preference in adults. Each two of the four colors were combined as a group to color the CPP tank (six groups in total). The 10-adult zebrafish with equal numbers of males and females (a shoal) swam freely in the color preference apparatus. After 5-min adaption, the proportion of numbers stayed in each colored zone was recorded every 1 min for 30-min experiment. Unlike the color preference of shoals, the individual fish was recorded the proportion of time spent in each colored zone during 30-min experiment. The behavioral analyses were performed with the SMART 3.0 software (SMARTSUPER, Panlab, Spain).

### 2.5. Statistical analysis

All experimental results were expressed as the means ± standard error of the mean (SEM) and analyzed by an independent t-test using the SPSS statistics program. Significance was set at p < 0.05 for all the experiments.

## 3. Results

### 3.1. The innate color preference of shoals in zebrafish

To investigate color preference in shoals of zebrafish, a shoal was introduced and allowed to swim freely in color-enriched CPP tank. After 5-min acclimation to the treatments, the location of each zebrafish in each colored zone was counted every 1 min for 30 min total of video recording. As demonstrated in Fig. 1A-1C, R (80.0% ± 16.56%) was preferred over Y (20.0% ± 16.56%) (t9 = 5.73, p<0.001), and R (77.10% ± 21.92%) was preferred over G (22.90% ± 21.92%) (t_9_ = 3.91, p<0.01), and Y (67.27% ± 16.52%) was preferred over G (32.73% ± 16.52%) (t_9_ = 3.31, p<0.01), suggesting the order of color preference was R>Y>G. Between B and G colors, the adults showed a greater preference for B (79.01% ± 12.65%) over G (20.99% ± 12.65%) (t_9_ = 7.25, p<0.001) (Figure 1D). However, no distinct preference was observed between R, Y and B (Fig. S1, Table S1). Thus, the order of RYGB preference was R>Y>G, B>G.

**Fig. 1.**
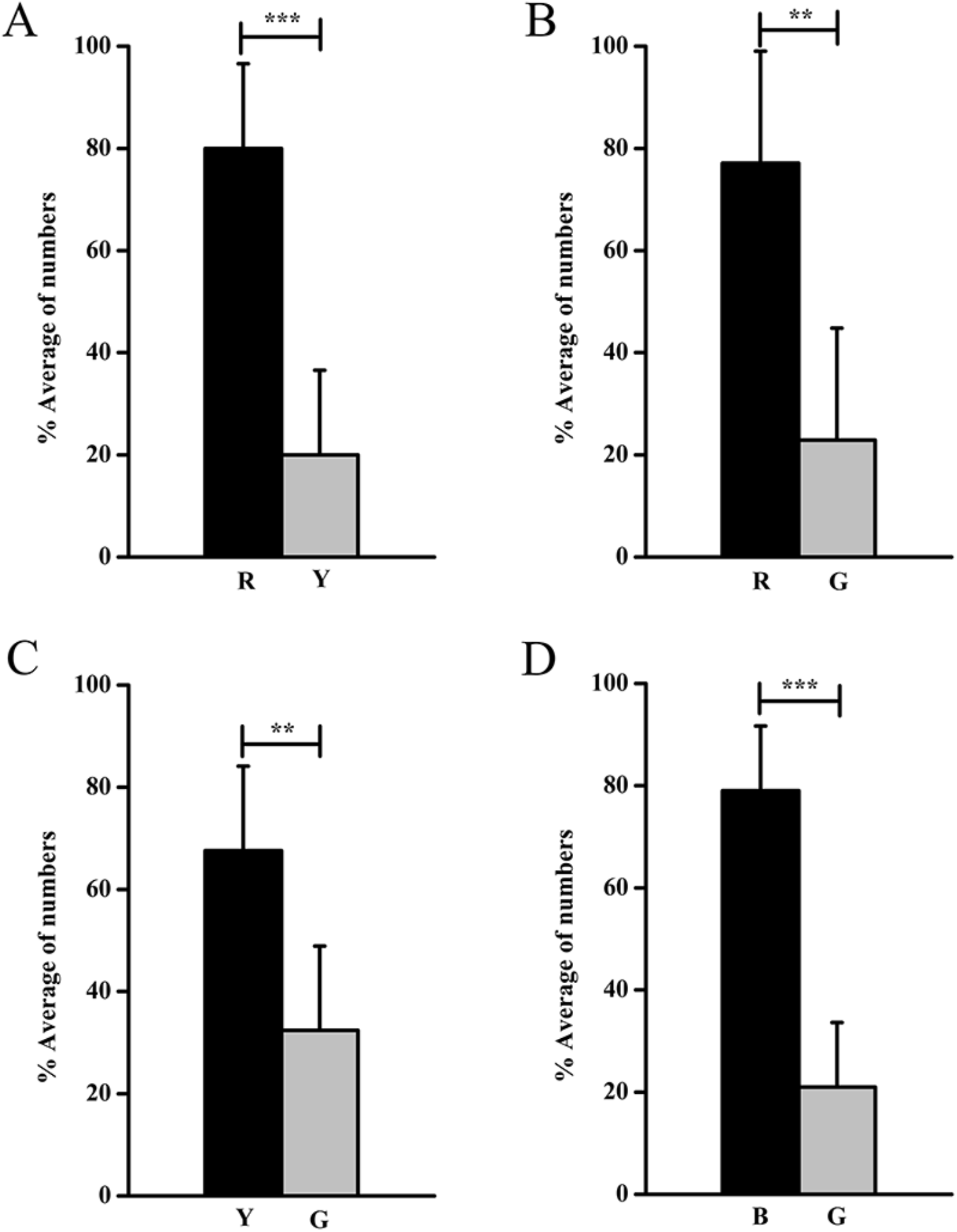
The shoals of zebrafish exhibit the innate color preference with 4 color combinations (R-Y, R-G, Y-G, B-G). R, red; Y, yellow; G, green; B, blue. The percentage of numbers of 10-adult zebrafish spent in each colored zone was counted every 1 min for a total of 30 min (n=10). ** p<0.01, *** p<0.001.

### 3.2. The innate color preference of a single zebrafish

We next set out to identify the social isolation that affected color preference in zebrafish. First, we focused on the color discrimination of an individual fish. To test the role of social contact in zebrafish, we measured color preference in an individual fish which was separated from a shoal. Individual fish could swim freely in the same CPP tank. After 5-min adaptation to the environment, the percentage of time spent in different zones was recorded during a period of 30 min. Unexpectedly, from Fig. 2 and Fig. S2, the zebrafish showed marked changes, completely losing their preference for all colors (R-Y, R-G, Y-G, B-G, B-R, B-Y) (Table S1). These data raised the possibility that social interaction in shoals of zebrafish played an important role in color preference.

**Fig. 2.**
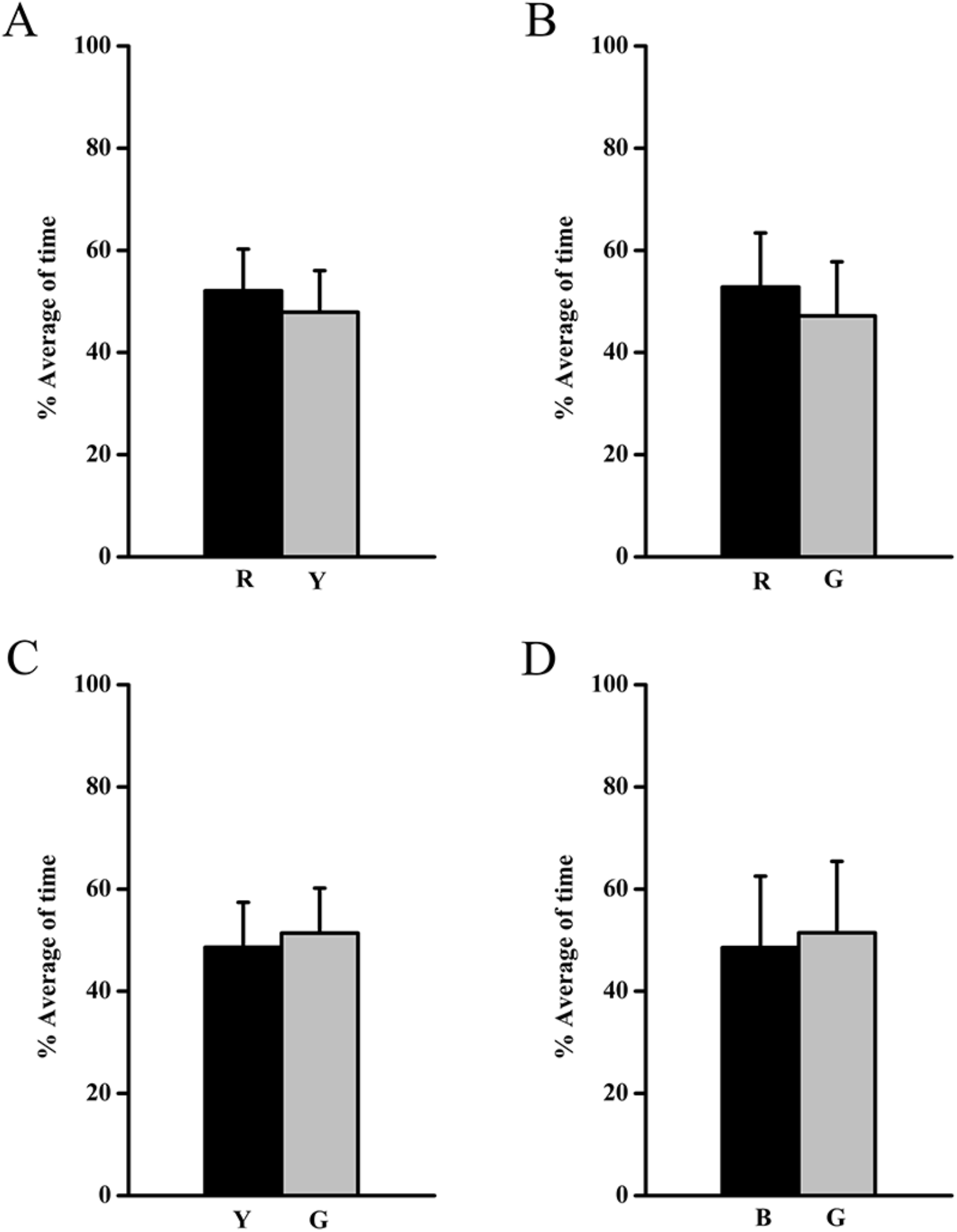
The color preference of an individual fish according to 4 color combinations (R-Y, R-G, Y-G, B-G). The percentage of time of fish spent in each colored zone was recorded during 30-min experiment (n=15).

### 3.3. The innate color preference of shoals depends on the social interaction

To start the color preference in shoals, a group of 10 fish with equal numbers of males and females remained in the D1-receptor antagonist solution for 30 min. Then, the shoals of zebrafish could swim freely in color-enriched CPP tank. After 5-min acclimation, the location of each zebrafish in each colored zone was counted every 1 min for 30 min total of video recording. From the results in Fig. 3 and Table S1, by contrast with the shoals without SCH23390 treatment, the shoals lost their interest in color combinations in color-enriched CPP tank (R-Y, R-G, Y-G, B-G). These results suggest that the color preference of shoals require the participation of social interaction.

**Fig. 3.**
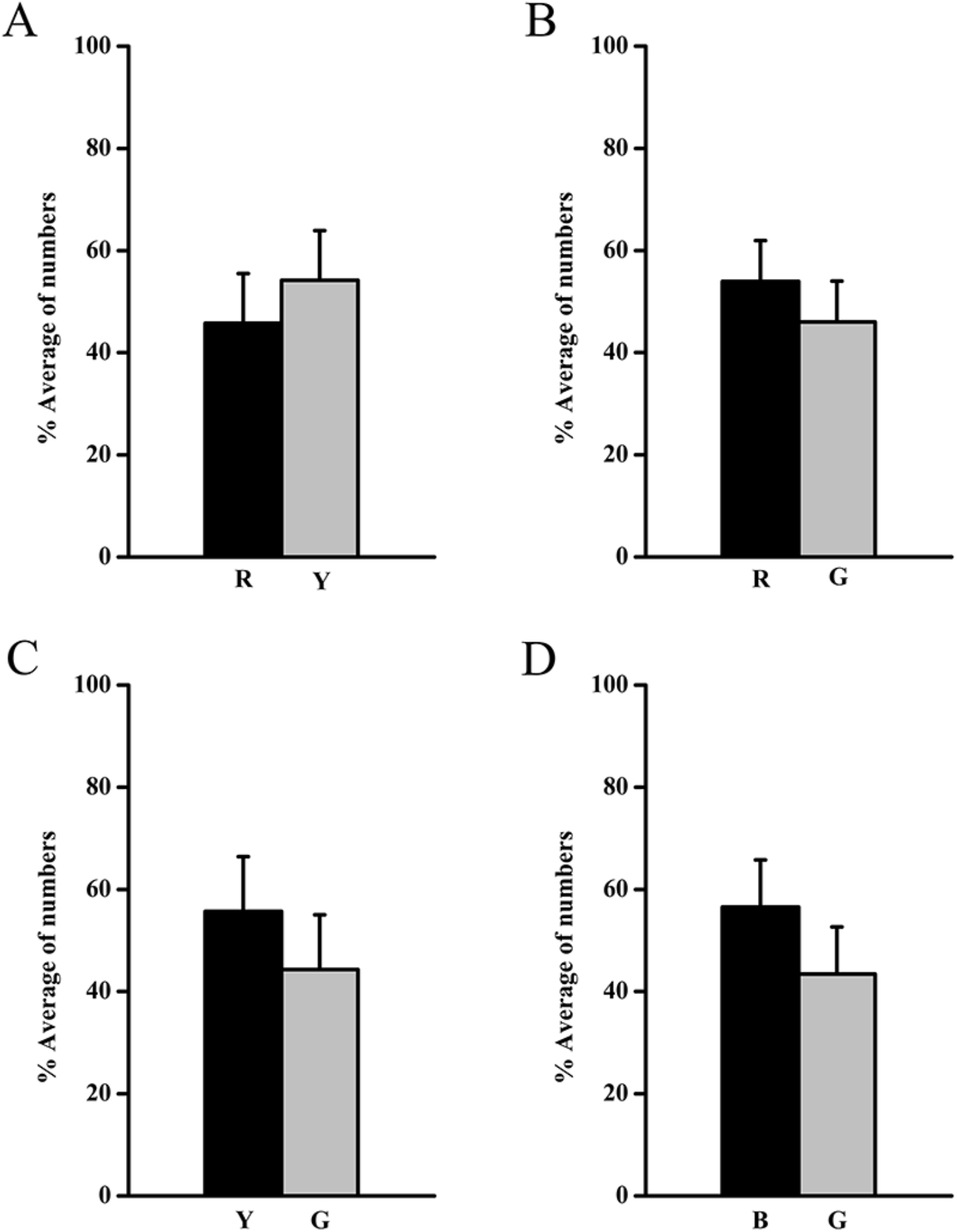
The color preference of shoals with SCH23390 treatment in 4 color combinations (R-Y, R-G, Y-G, B-G). The percentage of numbers of 10-adult zebrafish spent in each colored zone was counted every 1 min for a total of 30 min (n=10).

## 4. Discussion

The shoaling behavior is one of the most robust and consistent behavioral features in zebrafish, which has been observed both in nature [34] and in the laboratory [35]. Fish often forms aggregations and is found mostly in lakes, puddles, ponds, rice fields, ditches and small watercourses [36]. The intricate living conditions which contain abundant colors and shoaling behavior may affect foraging, predator avoidance and reproductive success. The results show a clear innate color preference in shoals of zebrafish. The innate color preference would spur shoal on to emigrate to the environment with a favorable color background that benefits social contact. According to this studies, the order of RYGB preference was R>Y>G, B>G. Some results of color preference were consistent with the findings which were reported by Park *et al*. [19] and Peeters *et al*. [37]. They used zebrafish larvae (5 dpf) to test the innate color preference in shoals, and found that zebrafish preferred R over Y, and R over G, and B over G. However, literature has emerged that offers contradictory findings about the innate color preference in shoals. A possible explanation for the contradictory conclusions are that the 5 dpf larvae did not develop shoaling behavior [20–22]. Therefore, we think that it is necessary to carry out controlled study which compares difference in color preference between individual and shoal adults.

So far, the color preference of individual fish has been studied by many groups, but no consensus is apparent with respect to this essential behavior in zebrafish. For example, Li *et al*. [38] found that zebrafish exhibited a robust preference for the green zone compared with the red zone. Soon after, an apparently contradictory result was reported by Pierog *et al*. [39], which demonstrated that zebrafish significantly preferred the red zone to the green zone. In addition, Kim *et al*. [40] claimed that there was no color preference between red and green. Since then, the color preference test has been replicated by different groups with their own experimental designs, it is difficult to control the experimental apparatus, the acclimation period and recording time. Park *et al*. [19] considered that vision of zebrafish can affect the color preference of fish by hypopigmentation of the retinal pigment epithelium.

Herein, for the first time, we identify the effect of social interaction that affects the color preference in shoal zebrafish. Our results demonstrated that individual zebrafish hardly had any clear preference for all the tested two-color combinations, while shoals of zebrafish exhibited a complex pattern of color preference and the order of RYGB preference was R>Y>G, B>G. We deduced that the color preference of zebrafish is an innate attribute of shoals, and therefore individual fish lack of social interaction with one another showed no color preference. As a social animal, zebrafish prefers to spend most of time to social group and develops a complex behavioral pattern depending on timely social contact with its companions [41–43]. Additionally, zebrafish exhibited a strong preference for their own phenotype, and such preference were mediated by visual signals [44]. Instead, socially deprived zebrafish failed to show social preference and social interaction [44,45]. These results suggested that social interaction could be responsible for eliciting the preference for colors.

The dopaminergic system is involved in several brain functions which has been found to be associated with the shoaling tendencies [46]. The dopamine plays key roles in the neurobehavioral functions in zebrafish [27,28]. For instance, the strong social stimuli will increase the dopamine and DOPAC levels in the brain of the adult zebrafish [27], and the short-term isolated zebrafish could reduce the level of DOPAC [28]. Dopamine receptors distribute in different brain regions. Clearly, the specific areas are involved in cognition, including hippocampus, the prefrontal cortex, the amygdale, and the ventral and dorsal parts of the striatum. There are four different dopamine receptor subtypes (D1, D2, D3, and D4) in the brain of zebrafish [30,47,48]. Among the different types of dopaminergic receptors, the excitatory D1 receptor (D1-R) subtype is the most predominately expressed in the brain regions [49]. D1-R activate the production of intracellular 3’-5’-cyclic adenosine monophosphate (cAMP) through adenylyl cyclase induction and regulate intercellular calcium signaling or protein kinase activity [50].

The social interaction in shoals can be affected by the dopaminergic system through influence of the dopamine level [29]. To investigate the role of social interaction in the innate color preference in shoals, we employed a D1 dopamine receptor antagonist (SCH23390) and analyzed its effects on behavior of color preference. The drug was chosen because D1-R antagonist, SCH23390, is most abundantly expressed dopamine receptor subtypes in the brain of zebrafish [30]. Second, SCH23390 disrupts social preference of zebrafish by decreasing the level of dopamine in dopaminergic system [31,32]. Finally, the drug is water soluble, and zebrafish can be administered by simple immersion in the drug solution.

The D1-R antagonist treatment which can disrupt the social interaction led to the deficits of color preference in shoals. The fish were employed by SCH23390 did not exhibit abnormal motor or posture and visual damage [29]. The article showed that SCH23390 decreased the preference of zebrafish to move toward and stay close to social stimuli [29]. Several potential mechanisms may be responsible for the decreased the level of dopamine through exposure to the D1 receptor antagonist (SCH23390). The decreased of dopamine levels imply the reduced dopamine production and/or increased dopamine degradation in response to the employed SCH23390 [51]. Yung *et al*. [52] have shown that D1-R localized in the post-synaptic terminals of neurons in the basal ganglia. The blockade of post-synaptic neurotransmitter receptors may impair signaling downstream and reduce neurotransmitter release. The antagonist of D1 receptor could increase the concentration of dopamine in the synaptic cleft which lead to reuptake and leakage to extra-synaptic areas. The increased extra-synaptic dopamine could activate dopaminergic autoreceptors on the pre-synaptic neuron and inhibit the dopamine synthesis [53]. Taken together, these studies and our own suggest the role of social interaction in innate color preference of shoals, with color discrimination deficits linked to the decreased dopamine level in the brain of zebrafish.

## Acknowledgements

We acknowledge the Funds for Huangshan Professorship of Hefei University of Technology (407-037019).

## Supplemental Information

**Fig. S1.**
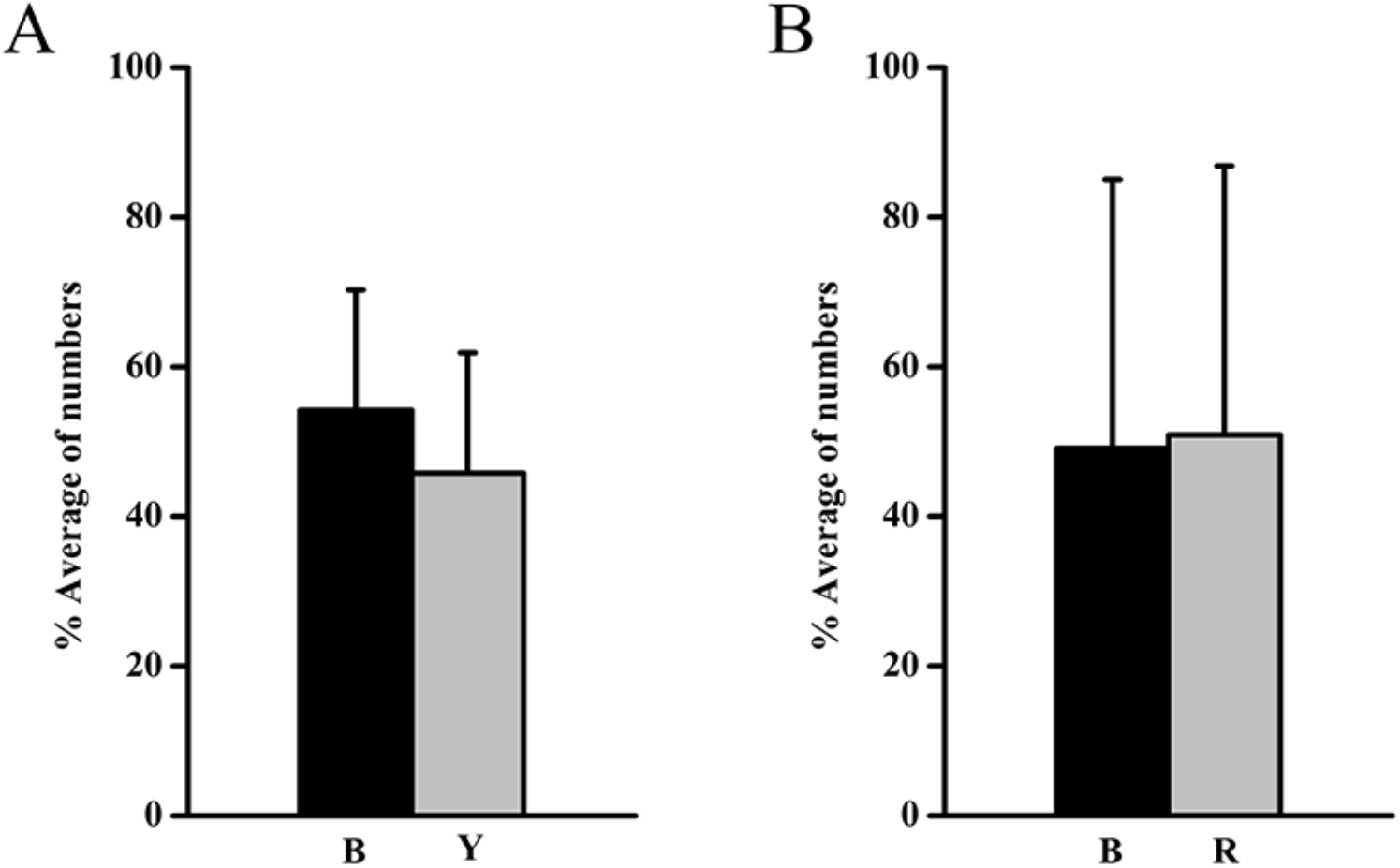
The color preference of shoals in B-Y and B-R. R, red; Y, yellow; B, blue. The percentage of numbers of 10-adult zebrafish spent in each colored zone was counted every 1 min for a total of 30 min (n=10).

**Fig. S2.**
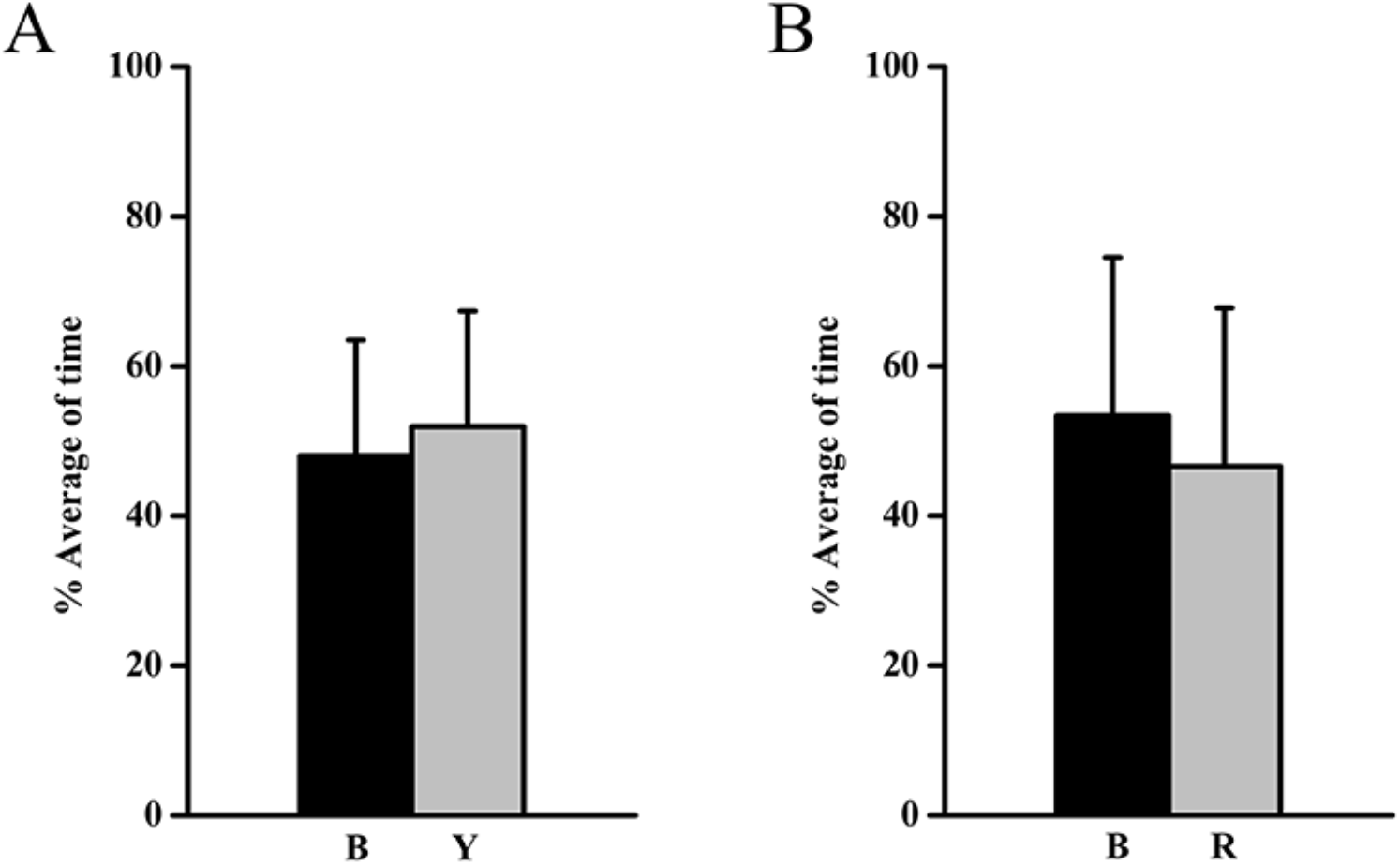
The color preference of an individual fish in B-Y and B-R. The percentage of time of fish spent in each colored zone was recorded during 30-min experiment (n=15).

**Table S1.** The results of color preference in zebrafish with different treatments.

| Treatment | Color | % Average of time (or % Average of number) | t-value, p-value |
| --- | --- | --- | --- |
| The color preference of shoals | R vs Y | 80.00±16.56 vs 20.00±16.56 | $t_9=5.73$, $p<0.001$*** |
| | R vs G | 77.10±21.92 vs 22.90±21.92 | $t_9=3.91$, $p<0.01$** |
| | Y vs G | 67.27±16.52 vs 32.73±16.52 | $t_9=3.31$, $p<0.01$** |
| | B vs G | 79.01±12.65 vs 20.99±12.65 | $t_9=7.25$, $p<0.001$*** |
| | B vs Y | 54.20±16.10 vs 45.80±16.10 | $t_9=0.83$, $p=0.431$ |
| | B vs R | 49.10±35.94 vs 50.90±35.94 | $t_9=0.08$, $p=0.939$ |
| The color preference of the single zebrafish | R vs Y | 52.10±8.14 vs 47.90±8.14 | $t_{14}=1.45$, $p=0.170$ |
| | R vs G | 52.82±10.59 vs 47.18±10.59 | $t_{14}=1.50$, $p=0.156$ |
| | Y vs G | 48.61±8.80 vs 51.39±8.80 | $t_{14}=0.88$, $p=0.393$ |
| | B vs G | 48.56±13.98 vs 51.44±13.98 | $t_{14}=0.57$, $p=0.578$ |
| | B vs Y | 48.07±15.42 vs 51.93±15.42 | $t_{14}=0.70$, $p=0.498$ |
| | B vs R | 53.38±21.15 vs 46.62±21.15 | $t_{14}=0.89$, $p=0.389$ |
| The color preference of shoals with SCH23390 treatment | R vs Y | 45.80±9.70 vs 54.20±9.70 | $t_9=1.37$, $p=0.204$ |
| | R vs G | 53.97±7.99 vs 46.03±7.99 | $t_9=1.57$, $p=0.151$ |
| | Y vs G | 55.70±10.72 vs 44.30±10.72 | $t_9=1.68$, $p=0.127$ |
| | B vs G | 56.57±9.20 vs 43.43±9.20 | $t_9=2.26$, $p=0.050$ |

## References

[1] P. Lucchetta, C. Bernstein, M. Théry, C. Lazzari, E. Desouhant, Foraging and associative learning of visual signals in a parasitic wasp, Anim. Cogn. 11 (2008) 525–533.

[2] J. Forrest, J.D. Tomson, Background complexity affects colour preference in bumblebees, Naturwissenschaften 96 (2009) 921–925.

[3] S. Lazopulo, A. Lazopulo, J.D. Baker, S. Syed, Daytime colour preference in Drosophila depends on the circadian clock and TRP channels, Nature 574 (2019) 108–111.

[4] Q. Duan, E. Goodale, R.C. Quan, Bird fruit preferences match the frequency of fruit colours in tropical Asia, Sci. Rep. 4 (2014) 5627.

[5] A. Avdesh, M.T. Martin-Iverson, A. Mondal, M. Chen, S. Askraba, N. Morgan, M. Lardelli, D.M. Groth, G. Verdile, R.N. Martins, Evaluation of color preference in zebrafish for learning and memory, J. Alzheimer’s Dis. 28 (2012) 459–469.

[6] Z.A. Bault, S.M. Peterson, J.L. Freeman, Directional and color preference in adult zebrafish: Implications in behavioral and learning assays in neurotoxicity studies, J. Appl. Toxicol. 35 (2015) 1502–1510.

[7] J. Oliveira, M. Silveira, D. Chacon, A. Luchiari, The zebrafish world of colors and shapes: Preference and discrimination, Zebrafish 12 (2015) 166–173.

[8] S.E. Palmer, K.B. Schloss, J. Sammartino, Visual aesthetics and human preference, Annu. Rev. Psychol. 64 (2013) 77–107.

[9] J.A. Endler, Signals, signal conditions, and the direction of evolution, Am. Nat. 139 (1992) S125–S153.

[10] A. Kelber, D. Osorio, From spectral information to animal colour vision: Experiments and concepts, Proc. R. Soc. B. 277 (2010) 1617–1625.

[11] P. Skorupski, L. Chittka, Is colour cognitive? Optics Laser Tech. 43 (2011) 251–260.

[12] K. Lunau, S. Papiorek, T. Eltz, M. Sazima, Avoidance of achromatic colours by bees provides a private niche for hummingbirds, J. Exp. Biol. 214 (2011) 1607–1612.

[13] J.A. Endler, D.A. Westcott, J.R. Madden, T. Robson, Animal visual systems and the evolution of color patterns: sensory processing illuminates signal evolution, Evolution. 59 (2005) 1795–1818.

[14] M.J. Hamblen-Coyle, D.A. Wheeler, J.E. Rutila, M. Rosbash, J.C. Hall, Behavior of period-altered rhythm mutants of Drosophila in light: dark cycles, J. Insect Behav. 5 (1992) 417–446.

[15] V.C. Fleisch, S.C. Neuhauss, Visual behavior in zebrafish, Zebrafish 3 (2006) 191–201.

[16] X. Li, B. Liu, X.L. Li, Y.X. Li, M.Z. Sun, D.Y. Chen, X. Zhao, X.Z. Feng, SiO2 nanoparticles change colour preference and cause Parkinson’s-like behaviour in zebrafish, Sci. Rep. 4 (2013) 1–9.

[17] R.M. Colwill, M.P. Raymond, L. Ferreira, H. Escudero, Visual discrimination learning in zebrafsh (Danio rerio), Behav. Process. 70 (2005) 19–31.

[18] N. Miller, R. Gerlai, Quantification of shoaling behaviour in zebrafish (Danio rerio), Behav. Brain Res. 184 (2007) 157–166.

[19] J.S. Park, J.H. Ryu, T.I. Choi, Y.K. Bae, S. Lee, H.J. Kang, C.H. Kim, Innate color preference of zebrafish and its use in behavioral analyse, Mol. Cells 39 (2016) 750–755.

[20] C. Buske, R. Gerlai, Shoaling develops with age in Zebrafish (Danio rerio), Prog. Neuro-Psychopharmacol. Biol. Psychiatry 35 (2011) 1409–1415.

[21] E. Dreosti, G. Lopes, A.R. Kampff, S.W. Wilson, Development of social behavior in young zebrafish, Front. Neural Circuits 9 (2015) 39.

[22] R.C. Hinz, G.G. de Polavieja, Ontogeny of collective behavior reveals a simple attraction rule, Proc. Natl. Acad. Sci. U. S. A. 114 (2017) 2295–2300.

[23] C. Buske, R. Gerlai, Maturation of shoaling behavior is accompanied by changes in the dopaminergic and serotoninergic systems in zebrafish, Dev. Psychobiol. 54 (2012) 28–35.

[24] P.M. Ma, M. Lopez, Consistency in the number of dopaminergic paraventricular organ-accompanying neurons in the posterior tuberculum of the zebrafish brain, Brain Res. 967 (2003) 267–272.

[25] J.A. Girault, P. Greengard, The neurobiology of dopamine signaling, Arch. Neurol. 61 (2004) 641–644.

[26] A. Vidal-Gadea, S. Topper, L. Young, A. Crisp, L. Kressin, E. Elbel, T. Maples, M. Brauner, K. Erbguth, A. Axelrod, A. Gottschalk, D. Siegel, J.T. Pierce-Shimomura, Caenorhabditis elegans selects distinct crawling and swimming gaits via dopamine and serotonin, Proc. Natl. Acad. Sci. U. S. A. 108 (2011) 17504–17509.

[27] M. Saif, D. Chatterjee, C. Buske, R. Gerlai, Sight of conspecific images induces changes in neurochemistry in zebrafish, Behav. Brain Res. 243 (2013) 294–299.

[28] S. Shams, S. Amlani, C. Buske, D. Chatterjee, R. Gerlai, Developmental social isolation affects adult behavior, social interaction, and dopamine metabolite levels in zebrafish, Dev. Psychobiol. 60 (2018) 43–56.

[29] T. Scerbina, D. Chatterjee, R. Gerlai, Dopamine receptor antagonism disrupts social preference in zebrafish: A strain comparison study, Amino. Acids. 43 (2012) 2059–2072.

[30] P. Li, S. Shah, L. Huang, A.L. Carr, Y. Gao, C. Thisse, B. Thisse, L. Li, Cloning and spatial and temporal expression of the zebrafish dopamine D1 receptor, Dev. Dyn. 236 (2007) 1339–1346.

[31] J.D. Steketee, Injection of SCH 23390 into the ventral tegmental area blocks the development of neurochemical but not behavioral sensitization to cocaine, Behav. Pharmacol. 9 (1998) 69–76.

[32] K. Kurata, R. Shibata, Effects of D1 and D2 antagonists on the transient increase of dopamine release by dopamine agonists by means of brain dialysis, Neurosci. Lett. 133 (1991) 77–80.

[33] J. Wang, C.H. Liu, F. Ma, W. Chen, J. Liu, B. Hu, L. Zheng, Circadian clock mediates light/dark preference in zebrafish *(Danio Rerio)*, Zebrafish 11 (2014) 115–121.

[34] R. E. Engeszer, L.B. Patterson, A.A. Rao, D.M. Parichy, Zebrafish in the wild: a review of natural history and new notes from the field, Zebrafish 4 (2007) 21–40.

[35] R. Gerlai, Social behavior of zebrafish: From synthetic images to biological mechanisms of shoaling, J. Neurosci. Methods 234 (2014) 59–65.

[36] R. Spence, M.K. Fatema, S. Ellis, Z.F. Ahmed, C. Smith, Diet, growth and recruitment of wild zebrafish in Bangladesh, J. Fish Biol. 71 (2007) 304–309.

[37] B.W.M.M. Peeters, M. Moeskops, A.R.J. Veenvliet, Color preference in Danio rerio: effects of age and anxiolytic treatments, Zebrafish 13 (2016) 330–334.

[38] X. Li, M.Z. Sun, X. Li, S.H. Zhang, L.T. Dai, X.Y. Liu, X. Zhao, D.Y. Chen, X.Z. Feng, Impact of low-dose chronic exposure to Bisphenol A (BPA) on adult male zebrafish adaption to the environmental complexity: Disturbing the color preference patterns and reliving the anxiety behavior, Chemosphere 186 (2017) 295–304.

[39] M. Pieróg, L. Guz, U. Doboszewska, E. Poleszak, P. Wlaź, Effect of alprazolam treatment on anxiety-like behavior induced by color stimulation in adult zebrafish, Prog. Neuropsychopharmacol. Biol. Psychiatry. 82 (2018) 297–306.

[40] Y.H. Kim, K.S. Lee, A.R. Park, T.J. Min, Adding preferred color to a conventional reward method improves the memory of zebrafish in the T-maze behavior model, Anim. cells and syst. 21 (2017) 374–381.

[41] N. Ruhl, S.P. McRobert, The effect of sex and shoal size on shoaling behaviour in Danio rerio, J Fish Biol. 67 (2005) 1318–1326.

[42] A. Etinger, J. Lebron, B.G. Palestis, Sex-assortative shoaling in zebrafish (Danio rerio), Bios. 80 (2009) 153–158.

[43] J.L. Snekser, N. Ruhl, K. Bauer, S.P. McRobert, The influence of sex and phenotype on shoaling decisions in zebrafish, Intern. J. Comp. Psychol. 23 (2010) 70–81.

[44] R.E. Engeszer, M.J. Ryan, D.M. Parichy, Learned social preference in zebrafish, Curr. Boil. 14 (2004) 881–884.

[45] G. Gerlach, A. Hodgins-Davis, C. Avolio, C. Schunter, Kin recognition in zebrafish: A 24-hour window for olfactory imprinting, Proc. Biol. Sci. 275 (2008) 2165–2170.

[46] P.S. Suriyampola, D.S. Shelton, R. Shukla, T. Roy, A. Bhat, E.P. Martins, Zebrafish social behavior in the wild, Zebrafish 13 (2016) 1–8.

[47] W. Boehmler, S. Obrecht-Pflumio, V. Canfield, C. Thisse, B. Thisse, R. Levenson, Evolution and expression of D2 and D3 dopamine receptor genes in zebrafish, Dev. Dyn. 230 (2004) 481–493.

[48] W. Boehmler, T. Carr, C. Thisse, B. Thisse, V.A. Canfield, R. Levenson, D4 Dopamine receptor genes of zebrafish and effects of the antipsychotic clozapine on larval swimming behavior, Genes Brain Behav. 6 (2007) 155–166.

[49] R.T. Fremeau, G.E. Jr, Duncan, M.G. Fornaretto, A. Dearry, J.A. Gingrich, G.R. Breese, M.G. Caron, Localization of D1 dopamine receptor mRNA in brain supports a role in cognitive, affective, and neuroendocrine aspects of dopaminergic neurotransmission, Proc. Natl. Acad. Sci. U. S. A. 88 (1991) 3772–3776.

[50] M. Naderi, A. Jamwal, D. P. Chivers, S. Niyogi, Modulatory effects of dopamine receptors on associative learning performance in zebrafish (Danio rerio), Behav. Brain Res. 303 (2016) 109–119.

[51] L. Diop, E. Gottberg, R. Brière, L. Grondin, T.A. Reader, Distribution of dopamine D1 receptors in rat cortical areas, neostriatum, olfactory bulb and hippocampus in relation to endogenous dopamine contents, Synapse 2 (1988) 395–405.

[52] K.K.L. Yung, J.P. Bolam, A.D. Smith, S.M. Hersch, B.J. Ciliax, A.I. Levey, Immunocytochemical localization of D1 and D2 dopamine receptors in the basal ganglia of the rat: Light and electron microscopy, Neurosci. 65 (1995) 709–730.

[53] A.H. Tissari, M.S. Lillgäls, Reduction of dopamine synthesis inhibition by dopamine autoreceptor activation in striatal synaptosomes with in vivo reserpine administration, J. Neurochem. 61 (1993) 231–238.

